# Neural Entrainment in the Theta Band Predicts Groove Perception in Popular Music

**DOI:** 10.1101/2025.11.11.687596

**Authors:** Dominik Keller, Tim Rohe, Alexander Raake

## Abstract

When humans listen to popular music, they often feel ‘the groove’, which is a pleasurable urge to move along with music. Previous studies suggest that groove arises from the brain’s entrainment to rhythmical patterns of music in the delta-theta and beta bands, which enables the brain to predict temporal structures. However, this notion has only been tested using simplified or artificial acoustic stimuli and melodies, rather than naturalistic popular music. In this study, we combined electroencephalography (EEG) measurements with two measures of neural entrainment to test whether neural entrainment in the delta, theta, and beta bands predicts individuals’ groove ratings of musical and clapping stimuli, which served as a control. We presented nine pop music songs and twelve MIDI-based clapping rhythms, which participants rated for their groove perception on a 5-point scale. Our results show that music songs received very variable groove ratings while ratings of clapping stimuli showed less variability. Univariate analyses demonstrated that both music and clapping stimuli consistently entrained neural oscillations in the delta–theta bands, but not beta bands, as measured by intertrial phase coherence (ITPC) and stimulus-brain coherence (SBC). ITPC and SBC in the theta band weakly, but significantly correlated with groove ratings in fronto-central clusters. Decoding of groove ratings from multivariate spatiotemporal-spectral EEG patterns of entrainment with curve-fitting and machine-learning models predicted groove perception with high accuracy, up to 0.8 Pearson correlation, where combined patterns of inter-trial phase coherence appeared more informative than patterns of stimulus-brain coherence. Our results support the theory that the perception of groove is rooted in the brain’s ability to entrain to ongoing dynamic temporal structures of music in the theta band. They further show that multivariate decoding models can be leveraged to predict subjective groove online from neural entrainment with high accuracy and temporal resolution.

**Summary:**

- While participants (n = 30) rated their subjective groove perception of 9 popular songs and 12 MIDI clapping rhythms, we measured how their brains entrained to these musical stimuli using stimulus-brain coherence (SBC) and inter-trial phase coherence (ITPC) measures.
- Both stimulus types strongly entrained delta–theta (1–7 Hz) oscillations. In univariate analyses, also both stimulus types at fronto-central, beat-related frequencies significantly predicted groove ratings.
- Multivariate decoding using EEG entrainment patterns predicted individual groove ratings far better (up to Pearson *r* = 0.8), with combined ITPC patterns providing the most information.

## 1 Introduction

When humans listen to the rhythm of certain types of music, they naturally synchronize with it, enhancing their enjoyment: they are in the groove [1]. In general, recurring beats, rhythms, and melodies in music naturally encourage movement, such as tapping, nodding, or dancing [2]. Madison define groove as “wanting to move some part of the body in relation to some aspect of the sound pattern” [2], while Janata et al. describe it as “an aspect of the music that induces a pleasant sense of wanting to move” [3]. These definitions, as well as a recent review [4], demonstrate that groove is a multidimensional psychological state which comprises perceptual, motivational, affective, and motor dimensions, because the state arises from the interaction of rhythm perception, sensorimotor prediction, and reward-related brain activity. While some studies distinguish between *feeling the groove, feeling in the groove*, and actively moving to music [3, 5, 6], this study primarily focuses on the experience of the overall *feeling the groove*, which we term groove in the remainder of the paper.

In general, the groove arises from rhythmical features of music such as musical isochrony, predictive timing [7], and binary subdivisions. A prototypical example for groovy music is swing, where swing notes emphasize the beat and syncopation [2]. However, previous research showed that groove does not only evolve from rhythmical features alone, but it requires a coupling of the human sensory and motor systems. For example, Janata et al. demonstrated that music with high ratings for ‘*being in the groove*’ elicits spontaneous rhythmic movements, and the quality of the sensorimotor coupling during a tapping task was inversely related to the experienced groove [3]. Other work showed that groove was correlated with low- and high-level music descriptors (see, e.g., [2, 8, 9]): Low-level factors such as the root mean square (RMS) curve, a measure of power, and low-frequency spectral flux, a measure of change of power, as well as event density, have proven effective predictors for groove ratings [3, 6, 10]. High-level musical factors can also influence groove: Subtle temporal variations in microtiming within a generally metronomic rhythm, such as syncopation and varying note durations, create a sense of groove, enjoyment, and pleasure [1, 11, 12, 7, 13], even though this finding is disputed [2, 9, 14]. Further, a higher number of instruments [5], bass drum patterns [15], low-frequency bass instruments [6], percussiveness and pulse clarity [10, 16] as well as beat saliency [17] promote stronger sensorimotor coupling, as found in clapping or whole-body movements, and groove ratings.

While a large body of research has focused on the musical factors that create groove, current research focused on the question of how the brain elicits the subjective feelings of groove. Neural oscillations, which indicate synchronization and information flow within and between neural networks[18, 19], are a prime candidate for a neural mechanism of groove. Neural oscillations entrain to external rhythmic inputs [20], for example to speech [21, 22] and music [23, 24, 25] in the delta-theta band (i.e., 1-7 Hz). This neural entrainment to the temporal envelope of a stimulus enables the brain to temporally predict and better process upcoming rhythmic stimuli, for example for parsing of speech segments [26], or to predict the multiple hierarchical regularities of music [27]. According to the neural resonance theory of music [28], neural oscillations entrain to musical events across multiple timescales so that the brain can form expectations of a multitude of temporal musical patterns. However, it is still unclear whether neural entrainment to naturalistic stimuli such as popular music is responsible for the feeling of groove. A key idea from the predictive coding account of music [29, 27] is that groove arises from the optimal balance between the regularity of the music (e.g., from a regular rhythm) and the uncertainty of musical structures (e.g., from the degree of syncopation). In this case, the brain is optimally challenged to predict rhythmic structures, resulting in an intermediate level of predictability (i.e., not too easy or too difficult) [30]. For example, the degree of syncopation relates to groove ratings with an inverse U-shaped relationship [7]). To resolve the prediction errors that result from partially uncertain predictions, the brain invokes motor planning to predict upcoming rhythmical structures [31], which we subjectively feel as the urge to move. Thus, previous studies have shown that rhythmic stimuli evoke neural entrainment not only in the delta-theta band, but also in the beta band (15–30 Hz), which arises from the coupling of auditory and motor networks in the cortex [23, 32, 33, 7, 34]. However, unlike our study, the mentioned ones did not use real music to evaluate groove.

However, whether neural entrainment to music mediates the individual feeling of groove has not been thoroughly tested with real music stimuli. Popular music songs, for example, elicit different strengths of the urge to move, with large inter-individual variability in groove sensation [3]. Further, previous studies investigated how the brain entrains to music [23, 24, 25] without investigating groove. Doelling et al., for example, recorded neural activity while both non-musicians and musicians listened to music at various tempi and performed a pitch-related task. The authors found that, for typical tempi, low-frequency entrainment in the delta-theta range (as measured by inter-trial phase coherence, or ITPC) reliably tracked musical stimuli in non-musicians, but not for slower rates. This entrainment correlated with task performance. In musicians, entrainment was enhanced across all tempi, suggesting that musical training fosters neural tracking of rhythmic structure. Other studies explored how groove is linked to simple auditory stimuli, such as highly regular tones [35, 36, 37], clapping rhythms [38], drumbeats [39], or simple melodies [7]. Cameron et al., for example, used rhythms from Steve

Reich’s *Clapping Music* to explore the relationship between neural entrainment and subjective feelings of groove. They compared human-performed rhythms, which naturally feature microtiming, with computer-programmed MIDI-based rhythms that are based on most precise timing. Measuring ITPC from EEG data, they found that neural entrainment in the delta band (1-4 Hz) correlated with behavioral groove ratings, but only for human-performed rhythms [38]. Most previous studies used ITPC as a measure of neural entrainment. ITPC quantifies whether a neural oscillation for the same stimulus elicits a coherent phase across stimulus repetitions in different trials [40]. Yet, neural entrainment to dynamic stimuli such as music can also be measured more directly for each music stimulus within a trial using stimulus–brain coherence (SBC). SBC is a spectral measure that quantifies the synchronization between the acoustic envelope of a continuous auditory stimulus and the electroencephalography (EEG) signal, capturing the similarity in the dynamics between the stimulus and the phase of neural oscillations [25, 41]. For example, Wollman et al. used SBC to show that theta (4-7 Hz) oscillations follow the acoustic envelope of music, and that this entrainment is modulated by melodic spectral complexity [25].

Overall, previous studies either investigated how the brain entrains to music without measuring subjective groove [23, 25, 24], or they investigate groove for simplified or artificial acoustic stimuli only [35, 36, 37, 38, 39, 7]. However, such regular acoustic stimuli may not convey the required intermediate level of regularity and unpredictability to arouse feelings of groove [27, 30, 29], as found in contemporary popular music. Thus, it remains unclear whether the groove sensation is linked to, or can be predicted by, neural entrainment to real musical stimuli, which can be measured by ITPC and SBC. In this study, we intend to address this issue in two ways: First, we aim to gain a deeper understanding of how neural entrainment to music contributes to the sensation of groove by analyzing ITPC and SBC, as suggested by predictive coding [27, 42]. Second, we aim to predict listeners’ groove sensation as accurately as possible from neural entrainment, thereby enabling an automated and time-resolved assessment of groove. Such models could generalize to new individuals and pop music stimuli and be used to quantify how properties of music or audio technology affect groove perception and overall music quality [4, 1, 43]. Therefore, in the present study, we combined univariate correlation and multivariate decoding analyses of neural entrainment, as measured by EEG, and groove ratings, to investigate how the brain entrains to simple clapping rhythms and pop songs. Further, we used state-of-the-art machine learning approaches to optimize the accuracy of predicting subjective groove on an individual level from neural entrainment patterns. For data collection, we conducted a subjective test in which participants were presented with 12 variants of clapping music used in [38] and 9 popular songs with a known range of groove ratings from low to high groove [3, 43].

Participants judged their subjective feeling of groove for each stimulus. To quantify neural entrainment to clapping rhythms and pop songs, we computed ITPC and SBC in the delta-theta band (1-7 Hz), which has been linked to prediction and prediction-error processing for rhythmic structures [27], and the beta band (15-30 Hz), which is associated with synchronization within sensorimotor networks [32, 33]. Because we presented looped replications of each song and clapping stimulus in each trial, ITPC and SBC could be compared and related to groove ratings at a single-trial level. To link neural entrainment to feelings of groove, we analyzed spatiotemporal-spectral entrainment patterns across multiple electrode channels, time points, and frequency bands within two frameworks: First, we computed the individual univariate correlations between the neural entrainment measures within the frequency bands and single channels of interest and the groove ratings. This approach allowed to investigate whether local neural entrainment mechanisms mediate groove. Second, we predicted trial-wise groove ratings from spatio-temporal-spectral patterns of neural entrainment with curve-fitting and machinelearning models. This approach enabled the exploitation of information from complex entrainment patterns of neural networks across channels, time points, and frequencies to predict groove. To prioritize generalizability, the models of the second approach were trained and crossvalidated (80-20 split) on data pooled across participants rather than as personalized models. This aims to report model prediction performance values that reflect the ability to predict groove for unseen individuals and stimuli rather than for a subject-specific fitting. Such predictive models can capture the multivariate structure of EEG entrainment signals and might enable real-time decoding and accurate prediction of groove. Furthermore, examining the most informative spatio-temporal-spectral features may generate more refined hypotheses about the neural computations driving groove.

## 2 Materials & Methods

In this section we describe participants, stimuli, experimental design and setup, EEG procedures (acquisition, pre-processing, and ITPC and SBC analyses), and the analyses of entrainment and groove ratings.

### 2.1 Participants

Thirty participants (11 females, 19 males) were recruited at Technische Universität Ilmenau, Germany. The age of the participants ranged from 22 to 53 years, with a mean age of 29.6 years (SD = 6.4). None of the participants used hearing aids or had any reported hearing disabilities. We did not control for or assess the participants’ musical background or expertise. The study was approved by the ethics committee of Technische Universität Ilmenau on 09 May 2023.

### 2.2 Stimuli

The experiment was divided into two independent blocks, as we aimed at testing neural entrainment and groove sensation for simple clapping rhythms and for more complex popular music songs. The order of blocks was counterbalanced so that half of the participants began with the clapping stimuli and the other half with the song stimuli. One block presented pop music songs (see Tab. 1), while the second block presented excerpts from a contemporary classical music piece called *Clapping Music*.

**Table 1:** Groove ratings of songs and IDs used in the study, as presented in Janata et al. [3]. Tempo was calculated using the *madmom* Python package. Janata’s groove ratings are normalized from Study 1 [3], and our groove ratings are normalized from our Study 1 published in [44].

| Songs | Tempo [bpm] | Duration [s] | Groove Ratings |  |
| --- | --- | --- | --- | --- |
|  |  |  | from [3] | from [44] |
| Can't Let Go | 190 | 10.1 | 0.46 | 0.44 |
| Come Fly With Me | 138 | 13.9 | 0.68 | 0.49 |
| Funk That | 140 | 6.9 | 0.62 | 0.50 |
| In The Mood | 163 | 11.7 | 0.76 | 0.70 |
| Please | 167 | 11.5 | 0.66 | 0.64 |
| Space Oddity | 94 | 14.3 | 0.30 | - |
| Superstition | 99 | 9.7 | 0.86 | 0.75 |
| The Child Is Gone | 97 | 7.4 | 0.49 | 0.26 |
| We Are More | 115 | 8.4 | 0.58 | 0.41 |

#### 2.2.1 Songs

Nine musical excerpts were chosen from Janata et al. [3] based on groove ratings from the original study and preliminary ratings from our own research [43, 44] (see Tab. 1). Each excerpt was edited to allow for five consecutive repetitions, with cuts made at musically meaningful points, in general after 4 or 8 bars, to preserve the rhythmic, harmonic, and melodic integrity. Consecutive repetitions were chosen to enable ITPC computations across repetitions acting as an ensemble of trials which is needed to calculate the measure. As a result, the final stimuli ranged from 34.5 to 71.5 s, with duration differences reflecting variations in tempo and harmonic structure. Additionally, two extra songs, one with low groove ratings and one with high groove ratings in both Janata’s and our pre-study, were used for training purposes.

#### 2.2.2 Clapping

We also used the twelve musical excerpts from Steve Reich’s *Clapping Music* as it was used by Cameron et al. [38]. Very detailed information about these stimuli can be found in that study. Being classified into contemporary classical music, it features twelve unique rhythms created through shifted overlaying of one basic clapping pattern with itself. Each rhythm spans twelve metrical positions with single claps, simultaneous claps, or rests. Unlike Cameron et al., we only played the MIDI based *mechanical* version as they were freely usable. To enable computation of ITPC across repetitions, we repeated each of the twelve excerpts ten times resulting in a stimulus length of around 20 s. Two randomly picked versions were used for training.

### 2.3 Experimental Design

The experiment was designed to assess participants’ immediate and intuitive perception of groove while their EEG was recorded. Prior to the main experimental runs, participants completed a training session during which they familiarized themselves with both the testing interface and the acoustic stimuli. The main experiment consisted of three experimental runs for the clapping and music stimuli each. Therefore, all participants listened to and rated each stimulus three times. The order of the clapping and music stimuli was randomized within each run; however, each participant completed all runs for one of the stimuli types before proceeding to the other. Within each presented stimulus, the music excerpt was repeated five times (songs) or ten times (clapping) to enable ITPC computations. The order of stimuli was fully randomized. Between the runs and the change of stimuli short breaks were taken. In every run, participants listened to the musical excerpts and then immediately rated their subjective experience of groove using a 5-point scale (1 = not groovy, 5 = very groovy) via a keyboard response. Ratings were subsequently rescaled to the range between 0 and 1 for modeling reasons. Participants were explicitly instructed to provide their ratings quickly and intuitively, without engaging in prolonged deliberation. To ensure consistency in data acquisition and to minimize ocular artifacts, participants maintained their gaze on a centrally presented fixation cross throughout each stimulus presentation and were advised to blink as little as possible during each trial. Further, to avoid motion artifacts in EEG data, participants were instructed to move as little as possible. As already pointed out, groove was operationally defined as “the aspect of music that induces a pleasant sense of wanting to move along with the music” [3].

### 2.4 Experimental Setup

The EEG data were recorded using a g.tec g.Nautilus Research EEG system ^1^ equipped with 32 wet/gel-based electrodes arranged according to the international 10-20 system with three different cap sizes to accommodate varying head circumferences. Data acquisition was performed on a computer with an i7-1185G7 processor running at 3 GHz, 64 GB of RAM, and a 2 TB SSD, operating under Ubuntu 22.04 with MATLAB R2022B Update 2. Participants were seated comfortably in a chair in front of a computer screen and two Geithain MO-2 speakers arranged in an equilateral triangle stereo setup, with a distance of 2 m from each other. The study took place in a distraction-free, neutrally colored, acoustically damped room (gray wall in front; gray curtains on the remaining walls) to minimize reverberation and maintain controlled listening conditions. During data collection, participants were instructed to remain still and maintain their gaze on a centrally positioned fixation cross (a + symbol) displayed on an otherwise black screen. Fig. 1 shows the experimental setup, which ensured both optimal EEG recording conditions and controlled presentation of the acoustic stimuli.

**Figure 1.**
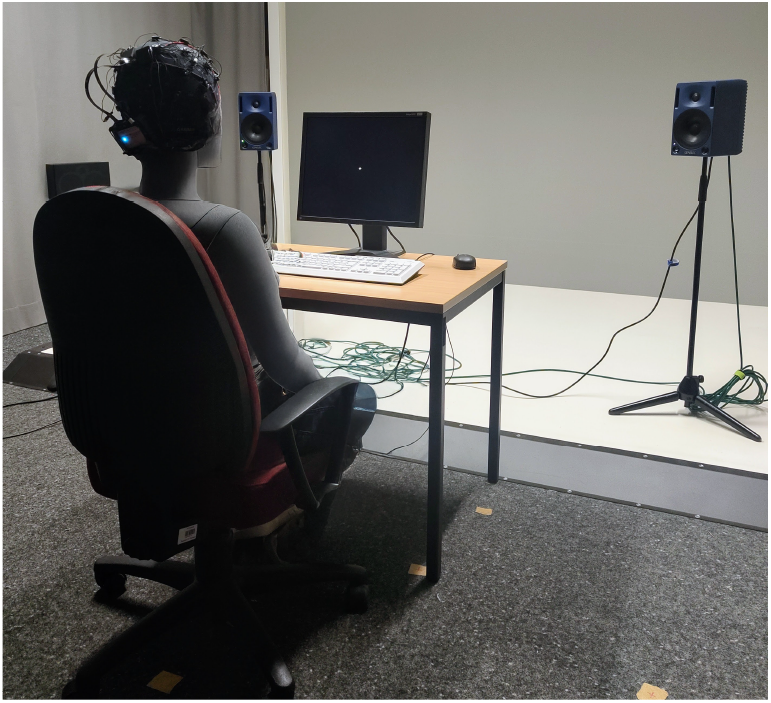
Experimental setup in the lab with a non-identifiable mannequin that is used to illustrate the participant position while fixating the central + symbol on the screen, with stereo loudspeakers reproducing the auditory stimuli. No actual participant is depicted in this figure.

### 2.5 EEG Data Acquisition and Preprocessing

EEG signals were recorded from 32 wet/gel-based electrodes positioned in an extended 10–20 montage using electrode caps and 32 channel DC amplifiers (g.tec, Nautilus Research, Austria). Electrodes were referenced to **FCz** using **AFz** as ground during recording. Signals were digitized at **500 Hz with a high-pass filter of 0.1 Hz**. Electrode impedances were kept below 30 kOhm. Eye blinks were automatically detected using data from the FP1 electrode. A blink was detected if the band-pass (1.5–15 Hz) filtered EEG signal exceeded two times the standard deviation, and the minimum duration between two consecutive blinks was 800 ms. Signal-space projectors (SSPs) were created from band-pass filtered (1.5–15 Hz) 400 ms segments centered on detected blinks, as implemented in Brainstorm [45]. The first spatial component of the SSPs was then used to correct blink artifacts in continuous EEG data. Further, all data were visually inspected for artifacts from blinks (i.e., residual blink artifacts after correction using SSPs), saccades, motion, electrode drifts or jumps. We had only a comparably low number of trials and a large proportion of trials was affected by artifacts because of their long duration (on average 55 % *±* 4 % standard error of the mean (SEM) for songs and 41 % *±* 4 % SEM for clapping), in contrast to typical EEG studies with a high number of trials with short duration. Thus, we decided to analyze all trials regardless of artifacts, assuming that they were not correlated to stimulus type or groove.

### 2.6 Analysis of Inter-Trial Phase Coherence and Stimulus-Brain Coherence

To compute inter-trial phase coherence (ITPC) and stimulus-brain coherence (SBC), we preprocessed the music and clapping stimuli using a time-frequency analyses of EEG data [24, 25]: We band-pass filtered the continuous EEG data between 0.25 and 85 Hz with a 50 Hz notch filter and segmented the data into epochs that started 3 s before and ended 3 s after each stimulus, depending on the stimulus duration (see Tab. 1). Using complex Morlet wavelets as implemented in Brainstorm [45], we extracted the spectral power and phase of single-trial EEG data of epochs from 1 to 30 Hz with increasing frequency steps (i.e., 1 to 2 Hz with 0.2 Hz steps, 2-10 Hz in 0.5 Hz steps, 10 to 20 Hz in 1 Hz steps and 20 to 30 Hz in 2 Hz steps, see [23]). The wavelet cycles increased linearly from 3 to 7 cycles across frequencies [40] and we used a full width at half maximum (FWHM) of 0.5 for the Gaussian taper. We downsampled all time-frequency representations to a sampling rate of 50 Hz and analyzed them in central channels (i.e., electrodes Cz, CP1, CP2, P3, Pz, P4). Further, we analyzed frequency data in all 32 individual channels.

To measure neural entrainment by coherence of neural phase, we computed ITPC between 1 and 30 Hz for each time-frequency point. ITPC was computed as absolute length of the mean phase angle vector [40] computed across replications of the same music (5 replications) or clapping stimulus (10 replications) within a trial.

This yielded one estimate of ITPC in each trial in channel-time-frequency space. In our univariate analyses, ITPC values were averaged across time for each stimulus (i.e., to align with SBC data, see below), and then analyzed as a function of channel and frequency. Because neural entrainment for clapping and music has been reported in central and parietal channels [23, 25], we averaged ITPC in these regions (i.e., electrodes Cz, CP1, CPz, CP2, P3, Pz, P4) in initial confirmatory univariate analyses. Importantly, we kept the temporal dimension of ITPC in our multivariate analyses to exploit potential temporal patterns. Thus, we normalized the time dimension for songs which had various durations (Tab. 1). Normalization was performed by binning the temporal dimension into 250 bins for each song.

To compute SBC as a measure of phase coherence between stimulus envelope and neural phase during a trial [25, 24], acoustic stimuli were first filtered using a cochlear-type gamma filter bank with 128 channels between 180 and 2205 Hz (i.e., half of the sampling frequency of the stimuli), as implemented in Matlab’s auditory toolbox [46]. To obtain the stimulus envelope, the channels’ outputs were Hilbert-transformed, the absolute values of the analytic signal were summed across the output channels, low-pass filtered with a cutoff of 50 Hz, and downsampled to 150 Hz. The stimulus envelope was finally time-frequency filtered using Morlet wavelets exactly as described for EEG data (see above), and downsampled a second time to 50 Hz to align the sampling rate to EEG data after time-frequency analyses. To compute SBC for each channel and frequency, we computed the single-trial coherence between stimulus envelope and EEG data, that is, the phase distance between envelope and EEG averaged across time using the following formula [24]:

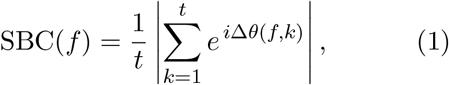

where Δ*θ*(*f, k*) is the angular distance between stimulus envelope and EEG at a single time point (k), with *t* time and *f* the considered frequency. We used only phase lags of zero between the EEG and envelope phases. Note that the SBC measure according to Equation (1) is formally independent from the amplitude of the EEG data. The computations yielded one SBC value per trial in channel-frequency space,. To test whether ITPC and SBC significantly deviated from chance level, we computed a baseline using a randomization strategy [24, 47]: For each stimulus, we randomly shifted phase data along the time dimension in each trial (n = 100 randomizations). Then we computed ITPC and SBC for randomized data as described above. Baseline-corrected ITPC and SBC was computed by subtracting the randomized ITPC or SBC, averaged over randomizations, from the original ITPC or SBC values.

To assess whether neural entrainment to music and clapping stimuli was significant in specific frequencies and channels, baseline-corrected ITPC and SBC were compared against zero across frequency-channels points at the group level. We used a non-parametric randomization test (5000 randomizations) in which we flipped the sign of the baseline-corrected ITPC or SBC [48] and computed one-sided one-sample t-tests as test statistic. To correct for multiple comparison across the frequency-channel points, we used a cluster-based correction [49] for p-values with the sum of the t-values across a cluster as cluster-level statistic and an auxiliary cluster-defining threshold of *t* = 2. To retain the spatial proximity of electrodes when forming clusters in the frequency-channel matrices, the electrode channels were ordered according to their one-dimensional proximity. Proximity was computed from the pairwise distance of electrodes in our montage using multidimensional scaling as implemented in Matlab.

### 2.7 Univariate Analysis of Entrainment and Groove Ratings

We correlated groove ratings with ITPC and SBC in univariate analyses to test whether the measures of neural entrainment in specific channels and frequencies predicted individual groove ratings. For this, we computed the Spearman correlations between per-trial and per-stimulus groove rating and ITPC and SBC features across all stimuli and trials (i.e., 9 x 3 songs or 12 x 3 clapping stimuli) for each participant and in each channel and frequency. To assess statistical significance of the correlations between groove ratings and neural entrainment measures at group level, we compared the individual correlations against zero using one-sample t-tests and cluster-based correction as described above for entrainment measures.

### 2.8 Multivariate Decoding and Modeling

In multivariate decoding analysis we modeled subjective groove ratings by predicting them from spatio-temporal-spectral entrainment patterns of ITPC and SBC using Python and scikitlearn [50]. This approach can enhance accuracy of the prediction of groove ratings compared to the univariate approaches by recognizing and utilizing the inherent multivariate structure of the patterns.

Spatio-temporal-spectral entrainment patterns were preprocessed as described above for univariate analyses, but we used the full complexity of the data dimensions to enhance prediction from complex patterns: In the spatial dimension, the patterns comprised all 32 channels. In the frequency dimension, the patterns comprised 0.3 up to 46 Hz (i.e., 48 frequency bands). For ITPC, the time dimension comprised 250 time bins after normalizing time across the songs or clapping stimuli. While SBC was computed within trials over time, ITPC was computed across trials, for each time bin. Thus, we pooled ITPC (across time and frequency) and also SBC (across frequency) using different functions. Pooling in this context refers to aggregating two- or multi-dimensional feature information into scalar (one-dimensional) summary features by using mathematic operations like mean, median, standard deviation, minimum, maximum, and several different percentile scores. This procedure produces a single scalar descriptor for the multidimensional data vectors consisting of electrode, frequency, and time information. This is done to reduce complexity, noise, and redundancy and to make decoding models more robust to input variations. We also aggregated ratings and pooled features in two ways that we call *averaged* and *individual. Averaged* indicates per-participant mean ratings (as each stimulus was played 3 times per participant), yielding one mean value each per participant–stimulus combination. For these averaged datasets, the SBC and ITPC features were averaged per participant. In contrast, *individual* marks per-participant, per-trial scores, so each participant contributed three ratings per stimulus, preserving trial-by-trial variability that is also available in the EEG measures. This approach increases sample size, potentially important for the multivariate decoding processes, and captures per-trial variability.

Our decoding strategy was implemented through a three-stage process characterized by progressively increasing complexity of the approaches using Python and scikit-learn [50]. In the initial stage, we employed a range of curvefitting parametric approaches including plain linear regression without regularization, Ridge regression, Lasso regression, and Tweedie regression. Subsequently, we applied simple and moderately complex machine learning techniques including random forest (RF) regressors, XGBoost (XGB), and support vector machines (SVM). They were each optimized by grid search within the 10-fold cross-validation framework on the training-validation split. We selected the models by testing various hyperparameter sets and chose the ones that minimized the mean squared error (MSE) on the validation folds. Finally, to be able to capture the highest level of complexity in the relationship between EEG data and groove ratings, we utilized AutoGluon’s *TabularPredictor*, a state-of-the-art AutoML framework that automates model selection, hyperparameter optimization, and ensembling [51]. In all the approaches, we trained the models using a randomized 10-fold cross-validation on 80 % of the data and then validated on the remaining 20 %. The best-performing model was selected based on minimizing MSE. We evaluated the model performances on the test set using RMSE (as this is in the same dimension as the ratings), *R*^2^, and Pearson correlation *r* to assess prediction accuracy and linear association with the ground truth.

We trained the models on six datasets for both the songs and clapping data. Those datasets consisted of the pooled SBC and ITPC data in different combinations and aggregations. For each subject, groove ratings are given per trial (individual) and are also averaged across trials that belong to the same stimulus (averaged). More details are provided in Tab. 2.

**Table 2:** Overview of data availability across conditions and feature extraction methods.

|  | Combined averaged | Combined individual | ITPC averaged | ITPC individual | SBC averaged | SBC individual |
| --- | --- | --- | --- | --- | --- | --- |
| ITPC & ratings per stimulus | x |  | x |  |  |  |
| ITPC & ratings per trial |  | x |  | x |  |  |
| SBC & ratings per stimulus | x |  |  |  | x |  |
| SBC & ratings per trial |  | x |  |  |  | x |

## 3 Results

We organize the results into three parts. Sec. 3.1 focuses on the behavioral groove ratings, including descriptive statistics and comparisons between the two stimulus sets to characterize differences in rating behavior. Sec. 3.2 presents analyses of neural entrainment (i.e., ITPC and SBC) to elucidate neural mechanisms of groove. Sec. 3.3 reports decoding analyses aimed at maximizing predictive performance for individual groove ratings using pooled features from ITPC, SBC, and their combination. We summarize model performance and feature contributions.

### 3.1 Behavioral Groove Ratings

The behavioral ratings showed that the song stimuli obtained differential groove ratings (main effect of song type, *F* (8, 801) = 57.86, *p* < 0.001), whereas the clapping stimuli led to much more similar ratings (main effect of clapping type, *F* (11, 1113) = 5.11, *p* < 0.001), with comparatively little variance between stimuli. The mean ratings incl. 95 % confidence intervals for both stimulus sets are shown in Fig. 2. For the *songs* stimuli the average ratings clearly varied, from low groove in *The Child Is Gone* and *Space Oddity* (0.29 and 0.30) to high groove in *Please* and *Superstition* (0.79 and 0.81). The small confidence intervals suggested consistent responses, highlighting a clear differentiation in perceived groove across songs. The between-participants similarity was very high with the mean correlation to the group mean resulting in 0.93 (*SD* = 0.05) and individual correlations ranging from 0.79 to 0.99. The within-participant Pearson correlation of 0.99 shows that the individuals were very consistent in their rating behavior during the test. The rating similarity with our pre-study showed high (*PCC* = 0.91) to moderate (Janata et al. [3], *PCC* = 0.67) agreement. The discrepancies can be likely attributed to differences in testing location (Germany and USA) and time (2012 and 2023), as well as other contextual or sociocultural factors that can affect subjective ratings. The mean ratings ranged from 0.29 to 0.81, covering about 52 % of the potential range, which suggested a broad variability in the groove ratings provided for songs.

**Figure 2.**
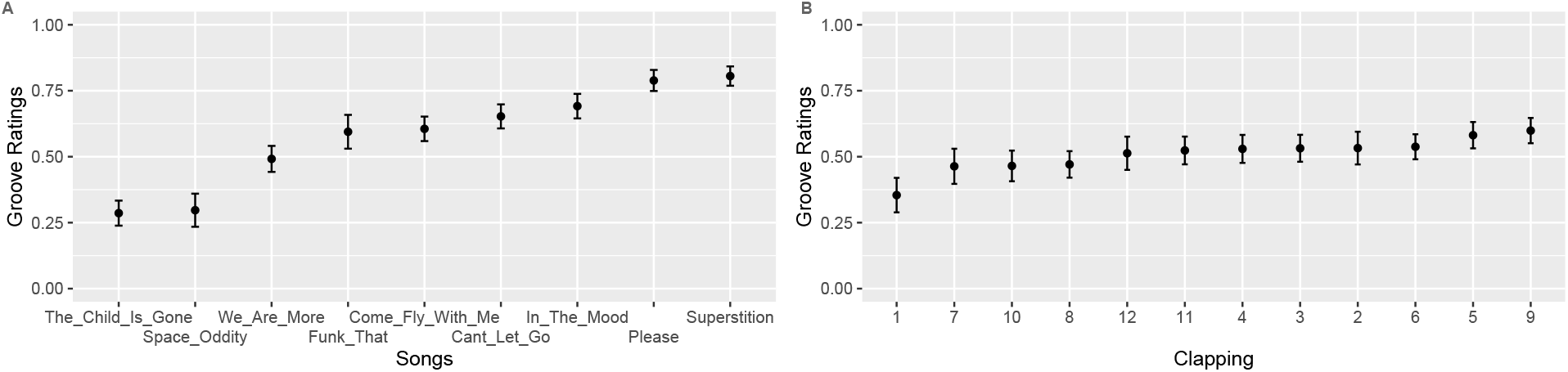
Mean and 95 % CIs of the normalized groove ratings for songs (A) and clapping (B).

In contrast, the *clapping* stimuli plot (see Fig. 2B) showed more narrowly clustered mean ratings, indicating less variation across stimuli, and a lower between-participants similarity. The mean Pearson correlation to the group mean was 0.69 (*SD* = 0.12), with a wider range from 0.41 to 0.92. The within-participants reliability, as measured by Pearson’s correlation, was 0.76. This indicated that there was more variability in individual ratings over time than for the *songs* stimuli. The rating similarity with Cameron et al.’s [38] average ratings was very low, showing a weak and negative relation of −0.20. As this paper looks into primarily into per-participant ratings, the difference in rating behavior is shortly addressed in Sec. 4. The overall mean ratings ranged from 0.35 to 0.60, covering only about 25 % of the potential range, which implied much less variation in the groove perception of clapping which is in line with the findings in [38].

### 3.2 Univariate Neural Entrainment

To understand the neural mechanisms of groove, we examined neural entrainment to songs and clapping using ITPC and SBC in central channels in the delta-theta and beta bands, for which neural entrainment for clapping and music have been reported [23, 38], and across all channels. Further, we tested how ITPC and SBC were related with behavioral groove ratings across trials.

#### 3.2.1 Inter-Trial Phase Coherence

First, we analyzed ITPC to songs and clapping stimuli across their repetitions within a trial in central channels. We found that ITPC was significantly larger than a randomization baseline in the delta and theta bands for most songs and clapping (one-sample t-tests against zero, *p* < 0.05; Fig. 3A). In contrast, ITPC was not significant in the beta band. The finding for the delta band in clapping music replicated the previous study that we built on here [38]. The profile of ITPC across frequencies showed distinct peaks in the delta and theta band at 1.3 Hz and 6 Hz for songs, and even narrower peaks at 1.7 Hz and 6 Hz for clapping. When we correlated the groove ratings with ITPC in individuals over different songs and clapping stimuli in the theta, delta, and beta bands (Fig. 3C), we found weak but significant correlations in the delta band for songs (mean *r* = 0.079, *t* = 2.157, *p* = 0.039, *Cohen*^*′*^*s d* = 0.394; one-sample ttests). For clapping, no correlations were significant (*p* > 0.05). Thus, we could confirm the results of no significant correlations between the delta ITPC and groove ratings for the clapping stimuli [38].

**Figure 3.**
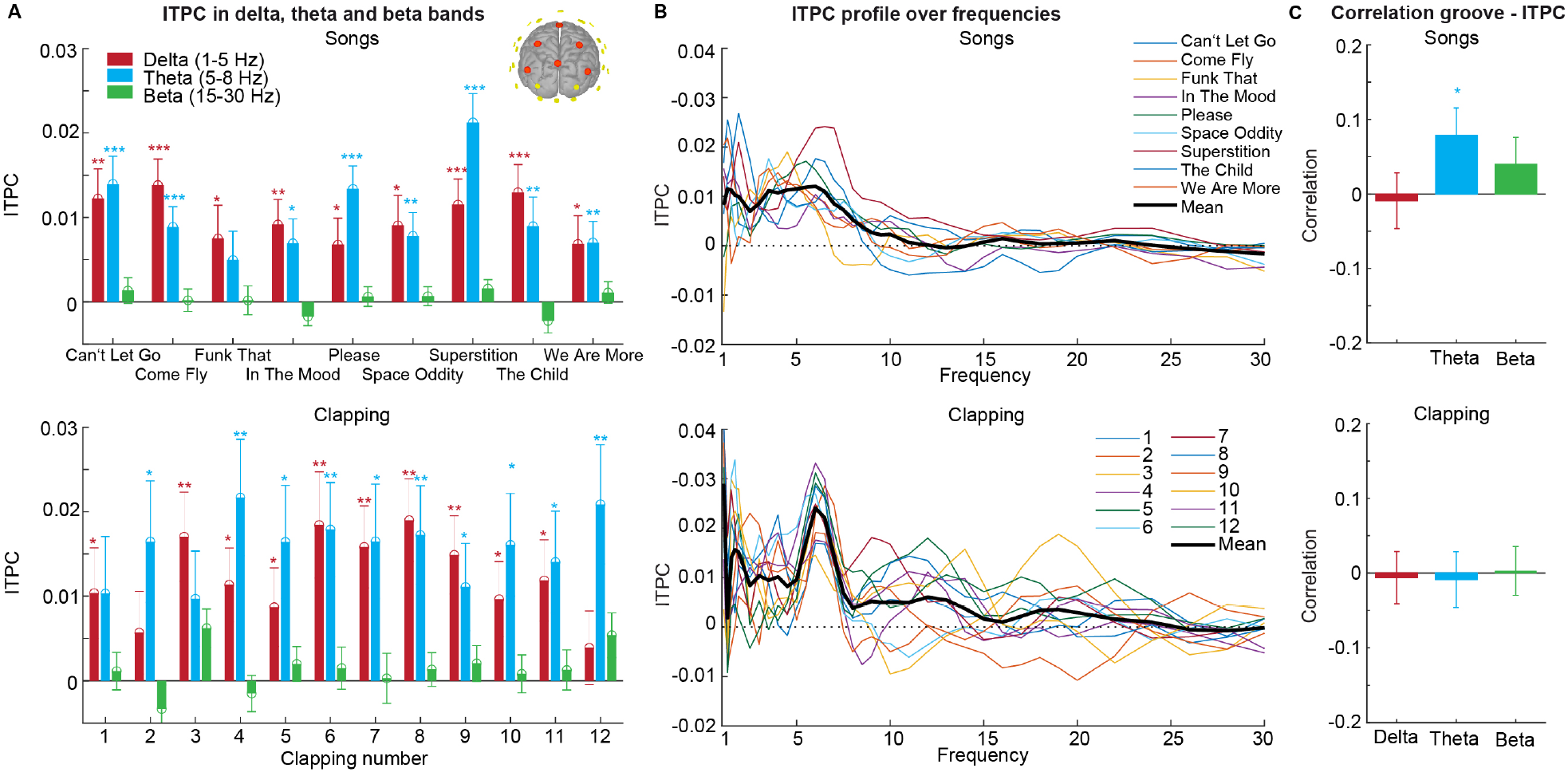
Inter-trial phase coherence (ITPC) and correlation between ITPC and groove ratings in the delta, theta and beta band in central channels, for songs (upper row) and clapping stimuli (lower row). (A) ITPC (*mean±SEM*) for the songs and clapping in the frequency bands compared to a randomization baseline with one-sample t-tests. p-values are multiple-comparison corrected across songs and clapping stimuli using Benjamini-Hochberg correction of the false-discovery rate. *** = *p* < 0.001, ** = *p* < 0.01, * = *p* < 0.05. (B) ITPC profile across the frequency spectrum. (C) Individual correlations (Fisher-z transformed; *mean ± SEM*) between ITPC and groove ratings. Individual correlations were tested against zero using one-sample t tests.

Next, we analyzed ITPC and its correlations with groove ratings across all channels (see Fig. 4). We found that all songs and clapping stimuli elicited strong ITPC clusters in the delta and theta bands using cluster-based randomization tests. For songs, the ITPC was strongest in two clusters in the fronto-central and frontal channels, and partially extended into the alpha band (Fig. 4A). For clapping, two similar frontal clusters emerged, and for some clapping stimuli ITPC was also significant in the beta band (Fig. 4C). The individual correlations between ITPC and groove ratings across songs or clapping stimuli revealed three significant cluster in the theta band, including the two clusters in fronto-central and frontal channels (Fig. 4B). The topography of the mean correlations showed that the correlations were weak (peak at *r ≈* 0.15) and peaked in frontocentral channels (Fig. 5A). Thus, higher theta-band ITPC was associated with higher groove ratings in songs. However, we did not find significant correlations for clapping stimuli (Fig. 4D).

**Figure 4.**
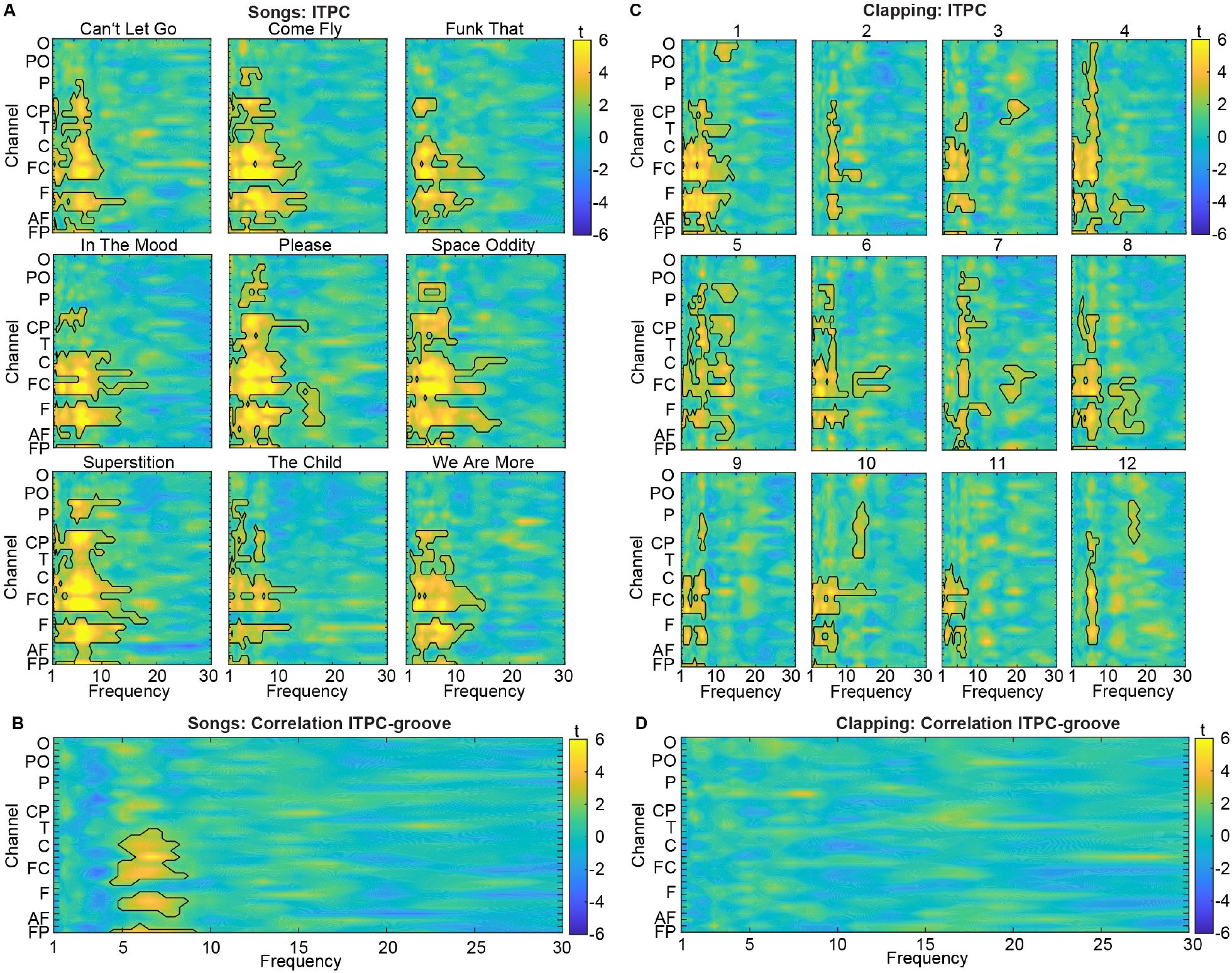
ITPC and correlation between ITPC and groove ratings between 1-30 Hz and in all 32 channels, for songs (A, B) and clapping (C, D). Frequency-channel plots show t-value maps for ITPC and correlations. Electrode channels are ordered according to their posterior-anterior proximity (O = occipital; PO = parietal-occipital; P = parietal; CP = central-parietal; T = temporal; C = central; FC = frontal central; F = frontal; AF = anterior-frontal; FP = fronto-polar). Significant clusters (*p* < 0.05; two-sided cluster-based corrected randomization t test) are demarcated by a solid line. (A) ITPC for the 9 songs compared to a randomization baseline. For song names A-I, see Table 1. (B) Individual correlations between ITPC and groove ratings. (C) ITPC for the 12 clapping rhythms compared to a randomization baseline. (D) Individual correlations between ITPC and groove ratings.

**Figure 5.**
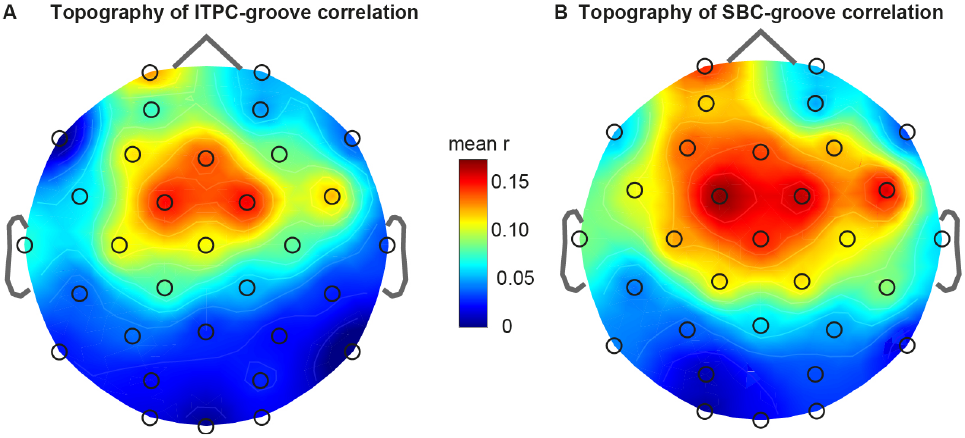
Topographies of mean individual correlation of neural entrainment to songs in the theta band (5.0 to 7.5 Hz) and groove ratings. (A) Correlation for ITPC. (B) Correlation for SBC.

Although the overall correlation pattern was weak, a more detailed exploratory analysis after pooling (for the pooling see Sec. 2.8) revealed that certain ITPC features showed localized, significant associations with groove for the song stimuli. See Tabs. 1 and 2 in the supplementary material for the highest-performing features and coefficients. Because the features were computed per stimulus and per trial, we correlated both the raw per-trial (individual) ratings and an aggregated per-stimulus (averaged) rating per subject with ITPC measures. This analysis identified three fronto-central clusters in the theta band where ITPC was significantly correlated with groove ratings; the strongest correlations were observed between about 5.5 and 7.5 Hz and localized to fronto-central electrodes such as FC1, FC2 and FP1. For example, the standard deviation of the ITPC in the FC1 channel at 6.5 Hz was significantly correlated with groove ratings (Pearson’s *r* = 0.316, *p* < 0.001), indicating that this particular ITPC-derived measure modestly predicted reported groove.

#### 3.2.2 Stimulus-Brain Coherence

When analyzing SBC for the songs and clapping stimuli within each trial and across all channels using cluster-based permutation tests (Fig. 6), we found significant clusters in six of the nine songs. In five songs (e.g., *Come Fly, In The Mood*, …), the clusters spread over the delta and theta bands in frontal and central channels. Only in the song *Funk That* a single significant cluster emerged in the beta band in central-parietal channels. For clapping, all twelve rhythms led to significant SBC clusters in the delta and theta bands in frontal and central channels. In seven clapping stimuli, SBC was also significant in the beta band, where clusters were distributed across frontal, central and parietal channels. Overall, the analyses of SBC showed significant entrainment to songs and clapping stimuli, most strongly in the delta and theta bands as previously reported in [25, 24]. In comparison to neural entrainment quantified by ITPC (Fig. 4), SBC manifested within similar clusters. However, ITPC revealed entrainment clusters with greater overall strength and statistical significance.

**Figure 6.**
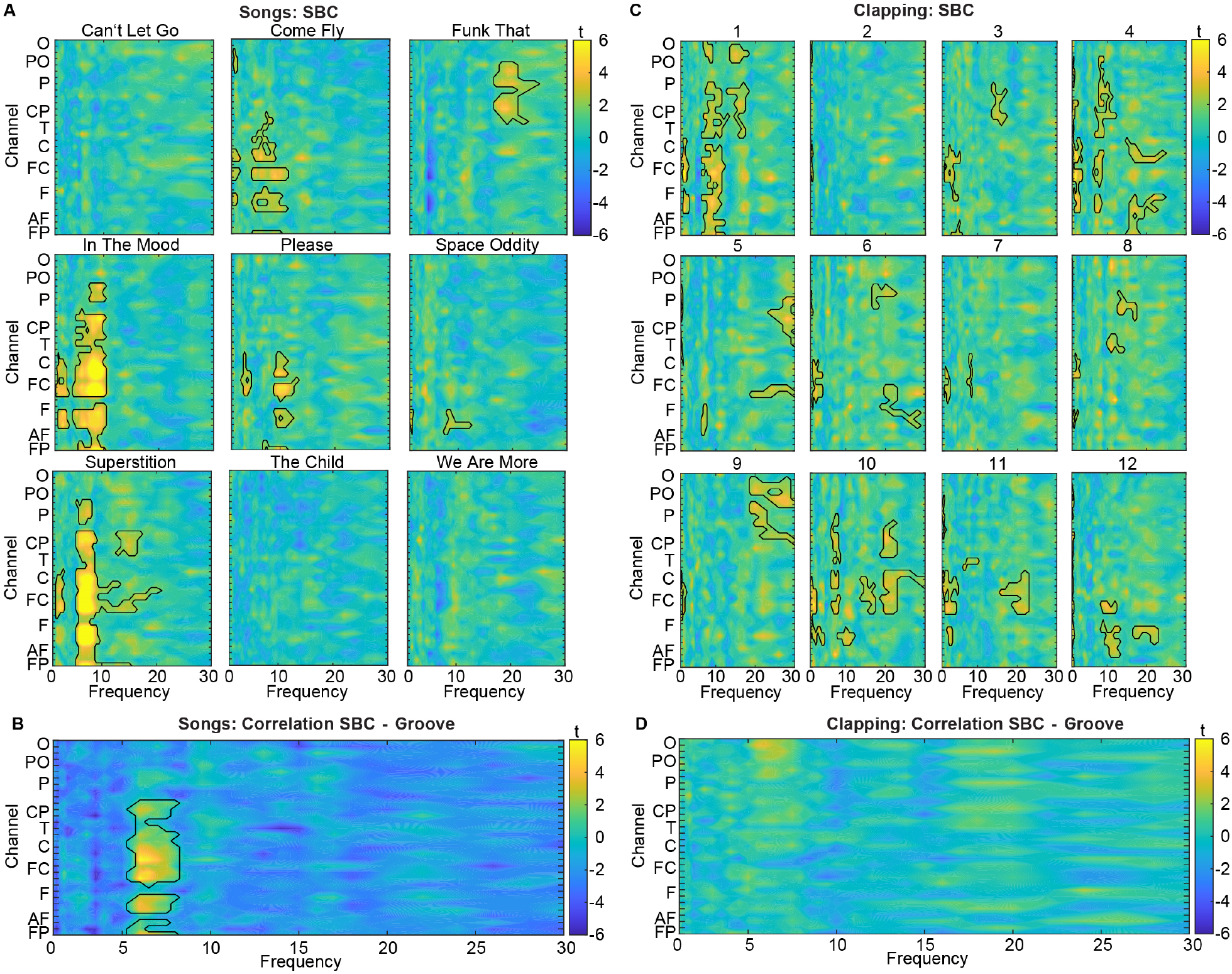
SBC and correlation between SBC and groove ratings between 1-30 Hz and all 32 channels, for songs (A, B) and clapping (C,D). Frequency-channel plots show t-value maps for SBC and correlations. Electrode channels are ordered according to their posterior-anterior proximity (O = occipital; PO = parietal-occipital; P = parietal; CP = central-parietal; T = temporal; C = central; FC = frontal central; F = frontal; AF = anterior-frontal; FP = fronto-polar). Significant clusters (*p* < 0.05; two-sided cluster-based corrected randomization t-test) are demarcated by a solid line. (A) SBC for the 9 songs compared to a randomization baseline. (B) Individual correlations between SBC and song groove ratings. (C) SBC for the 12 clapping stimuli compared to a randomization baseline. (D) Individual correlations between SBC and groove ratings of clapping.

When correlating individual SBC with individual groove ratings, we found that SBC significantly predicted groove ratings in three frontal and central clusters in the theta band with a peak around 6 Hz (Fig. 6C). Closer inspection of the topography of correlations showed that the correlations were weak (peak at *r ≈* 0.175) and peaked in fronto-central channels (Fig. 5B). The clusters and topography of significant correlations appeared remarkably similar to the correlations of ITPC. Similar to ITPC, we did not find significant clusters of correlations between SBC and groove ratings for clapping stimuli (Fig. 6D). To further analyze these relationships, we correlated both the raw per-trial (individual) ratings and the aggregated per-stimulus (averaged) rating per subject with pooled SBC features. This finer-grained exploratory analysis confirmed significant SBC–groove associations in the three frontal/central theta clusters and showed that the highest correlations concentrated between approximately 5.5 and 7.5 Hz in fronto-central electrodes such as FC1, FC2 and FP5. The strongest distinct SBC features and their Pearson and Spearman correlation coefficients are reported in Tabs. 1 and 2 in the supplementary material. Although these SBC–groove correlations for songs were modest in magnitude, the clapping stimuli again revealed no significant correlations; nonetheless, a number of individual SBC features for clapping showed significant positive or negative correlations that spanned different frequencies, electrodes and temporal windows.

To summarize, our univariate analyses revealed strong entrainment for ITPC and SBC in the delta-theta band in frontal and central channels for both songs and clapping, but only SBC was consistently correlated with subjective groove ratings for songs, albeit the correlation was modest on average.

### 3.3 Multivariate Decoding of Groove

Although univariate correlations between entrainment and groove perception were significant but low, information on groove may be encoded more widely in complex spatio-temporal-spectral patterns of entrainment. Thus, we next tested whether multivariate decoding approaches may learn to better predict groove ratings from such patterns 2.8. To predict groove ratings from the full feature space, we applied five decoding approaches, the parametric Ridge, the three traditional machine learning approaches RF, XGB, and SVM, and AutoGluon’s TabularPredictor. Models were trained on entrainment data from all participants rather than as personalized models using cross validation. Thus, the performance may reflect a generalization to unseen individuals and stimuli rather than subject-specific fits. Detailed decoding performance data are shown in Tab. 3 for the song datasets and in Tab. 4 for the clapping dataset.

**Table 3:** Average metrics (RMSE, *R*^2^, Pearson’s *r*, significance) per model and dataset for the songs stimuli. Significance codes: ns (*p* ≥ 0.05), * (*p* < 0.05), ** (*p* < 0.01), *** (*p* < 0.001).

| Model | Combined averaged |  |  |  | ITPC averaged |  |  |  | SBC averaged |  |  |  |
| --- | --- | --- | --- | --- | --- | --- | --- | --- | --- | --- | --- | --- |
| | RMSE | $R^2$ | $r$ | Sig. | RMSE | $R^2$ | $r$ | Sig. | RMSE | $R^2$ | $r$ | Sig. |
| Ridge | 0.272 | 0.069 | 0.297 | ns | 0.276 | 0.045 | 0.247 | ns | 0.270 | 0.089 | 0.328 | * |
| Random Forest | 0.280 | 0.017 | 0.221 | ns | 0.282 | 0.007 | 0.130 | ns | 0.265 | 0.122 | 0.442 | ** |
| XGBoost | 0.273 | 0.069 | 0.282 | ns | 0.290 | -0.053 | 0.001 | ns | 0.254 | 0.192 | 0.457 | ** |
| SVR | 0.276 | 0.047 | 0.271 | ns | 0.280 | 0.021 | 0.220 | ns | 0.269 | 0.096 | 0.337 | * |
| AutoGluon | 0.256 | 0.163 | 0.427 | *** | 0.284 | -0.033 | -0.008 | ns | 0.248 | 0.212 | 0.471 | *** |

Table 3: Average metrics (RMSE, $R^2$ , Pearson’s $r$ , significance) per model and dataset for the **songs** stimuli. Significance codes: ns ( $p \geq 0.05$ ), \* ( $p < 0.05$ ), \*\* ( $p < 0.01$ ), \*\*\* ( $p < 0.001$ ).
| Model | Combined individual |  |  |  | ITPC individual |  |  |  | SBC individual |  |  |  |
| --- | --- | --- | --- | --- | --- | --- | --- | --- | --- | --- | --- | --- |
| | RMSE | $R^2$ | $r$ | Sig. | RMSE | $R^2$ | $r$ | Sig. | RMSE | $R^2$ | $r$ | Sig. |
| Ridge | 0.153 | 0.727 | 0.854 | *** | 0.165 | 0.674 | 0.827 | *** | 0.281 | 0.075 | 0.293 | *** |
| Random Forest | 0.212 | 0.477 | 0.837 | *** | 0.214 | 0.454 | 0.806 | *** | 0.279 | 0.093 | 0.381 | *** |
| XGBoost | 0.167 | 0.673 | 0.842 | *** | 0.171 | 0.652 | 0.821 | *** | 0.260 | 0.209 | 0.474 | *** |
| SVR | 0.178 | 0.630 | 0.820 | *** | 0.186 | 0.589 | 0.789 | *** | 0.280 | 0.083 | 0.301 | *** |
| AutoGluon | 0.165 | 0.688 | 0.834 | *** | 0.162 | 0.700 | 0.837 | *** | 0.262 | 0.215 | 0.475 | *** |

**Table 4:** Average metrics (RMSE, *R*^2^, Pearson’s *r*, significance) per model and dataset for the **clapping** stimuli. Significance codes: ns (*p* ≥ 0.05), * (*p* < 0.05), ** (*p* < 0.01), *** (*p* < 0.001)

| Model | Combined averaged |  |  |  | ITPC averaged |  |  |  | SBC averaged |  |  |  |
| --- | --- | --- | --- | --- | --- | --- | --- | --- | --- | --- | --- | --- |
| | RMSE | $R^2$ | $r$ | Sig. | RMSE | $R^2$ | $r$ | Sig. | RMSE | $R^2$ | $r$ | Sig. |
| Ridge | 0.228 | -0.008 | 0.165 | ns | 0.232 | -0.045 | 0.095 | ns | 0.221 | 0.057 | 0.287 | * |
| Random Forest | 0.227 | 0.006 | 0.152 | ns | 0.229 | -0.010 | 0.061 | ns | 0.222 | 0.051 | 0.280 | ns |
| XGBoost | 0.228 | -0.006 | 0.136 | ns | 0.232 | -0.037 | 0.028 | ns | 0.224 | 0.026 | 0.218 | ns |
| SVR | 0.235 | -0.021 | 0.200 | ns | 0.229 | -0.016 | 0.044 | ns | 0.219 | 0.072 | 0.303 | * |
| AutoGluon | 0.213 | 0.054 | 0.240 | *** | 0.222 | -0.026 | 0.035 | ns | 0.208 | 0.099 | 0.317 | *** |

Table 4: Average metrics (RMSE, $R^2$ , Pearson’s $r$ , significance) per model and dataset for the **clapping** stimuli. Significance codes: ns ( $p \geq 0.05$ ), \* ( $p < 0.05$ ), \*\* ( $p < 0.01$ ), \*\*\* ( $p < 0.001$ )
| Model | Combined individual |  |  |  | ITPC individual |  |  |  | SBC individual |  |  |  |
| --- | --- | --- | --- | --- | --- | --- | --- | --- | --- | --- | --- | --- |
| | RMSE | $R^2$ | $r$ | Sig. | RMSE | $R^2$ | $r$ | Sig. | RMSE | $R^2$ | $r$ | Sig. |
| Ridge | 0.293 | -0.083 | 0.093 | ns | 0.298 | -0.117 | 0.047 | ns | 0.280 | 0.011 | 0.149 | ns |
| Random Forest | 0.281 | 0.003 | 0.107 | ns | 0.282 | -0.001 | 0.062 | ns | 0.278 | 0.028 | 0.198 | * |
| XGBoost | 0.280 | 0.007 | 0.137 | ns | 0.285 | -0.023 | 0.059 | ns | 0.283 | -0.014 | 0.116 | ns |
| SVR | 0.282 | -0.005 | 0.067 | ns | 0.282 | -0.005 | 0.067 | ns | 0.282 | -0.002 | 0.086 | ns |
| AutoGluon | 0.267 | 0.097 | 0.315 | *** | 0.267 | 0.097 | 0.315 | *** | 0.278 | 0.022 | 0.154 | *** |

We pre-evaluated plain linear regression without regularization, Ridge, Lasso, and Tweedie regressions. Linear and Lasso overfitted by having very high training performance and very low test correlations across all datasets, while Ridge and Tweedie performed similar and comparable between training and validation. For simplicity reasons, we therefore only report the Ridge regression results from this group of models. In the second stage, RF, XGB, and SVR regressors were optimized via grid search within the cross-validation framework. For the most complex modeling, AutoGluon’s TabularPredictor was used for automated model selection, optimization, and ensembling.

Different sets of features revealed the best performances for decoding groove for the songs and clapping stimuli. The test metrics RMSE, *R*^2^, as well as Pearson’s *r* and its significance varied between the stimuli and the different datasets, see the results in Tab. 2. Decoding performance for the clapping stimuli showed much weaker and mostly non-significant Pearson correlations. For both stimuli sets, better decoding performances were found on the per-subject per-trial (individual) data, with best performance across all models for the combined datasets. In case of the songs stimuli, Pearson’s *r* was on average > 0.85 across all models, with p-values indicating significance (***), which showed a good decoding performance. Ridge and XGB delivered the best overall fit on average, yielding the lowest RMSE and highest *R*^2^. SVR, despite tuning its penalty and kernel width, and AutoGluon exhibited still very comparable predictive performances. Importantly, for the individual songs datasets all five modeling approaches produced statistically significant high Pearson correlations between predicted and actual groove ratings, which was not the case for the clapping stimuli sets. There, only AutoGluon was able to consistently derive statistically significant models for the individual datasets. Together, these findings indicated a reliable, systematic relationship between the predictors and groove ratings that generalized across stimulus types.

However, from the averaged datasets, the SBC data proved to be more correlated to the groove ratings. Therefore, for the songs data, SBC data could be used to decode groove to when averaged and ITPC data produced better results for individual datasets.

To exemplarily showcase the robustness of the decoding results across cross-validation folds, Tab. 5 depicts more detailed performance data including mean values and standard deviations for the XGBoost approach and the songs datasets, allowing to assess how the results vary between cross-validation folds. This decoding approach was exemplarily chosen as it delivers the combination of high decoding performance while it is still relatively efficient to train. The performance data across all datasets were similar in terms of metrics like RMSE, Pearson’s *r*, and *R*^2^. This indicated that model accuracy was largely independent of the specific data points used for training and testing in different cross-validation folds. Notably, the best-performing datasets converged on highly similar performance metrics. This suggested that the predictive structure of neural entrainment generalized well across individuals and musical stimuli.

**Table 5:** Exemplary XGB decoding (*mean ± SD* across cross-validation) across songs datasets. Significance codes: ns (*p* ≥ 0.05), ** (*p* < 0.01), *** (*p* < 0.001).

| Dataset | RMSE | $R^2$ | Pearson's $r$ | Sign. |
| --- | --- | --- | --- | --- |
| combined averaged | $0.273 \pm 0.021$ | $0.069 \pm 0.046$ | $0.282 \pm 0.103$ | ns |
| combined individual | $0.167 \pm 0.011$ | $0.673 \pm 0.034$ | $0.842 \pm 0.019$ | *** |
| ITPC averaged | $0.290 \pm 0.017$ | $-0.053 \pm 0.069$ | $0.001 \pm 0.112$ | ns |
| ITPC individual | $0.171 \pm 0.010$ | $0.652 \pm 0.045$ | $0.821 \pm 0.032$ | *** |
| SBC averaged | $0.254 \pm 0.022$ | $0.192 \pm 0.049$ | $0.457 \pm 0.070$ | ** |
| SBC individual | $0.260 \pm 0.009$ | $0.209 \pm 0.052$ | $0.474 \pm 0.064$ | *** |

Overall, our decoding analyses demonstrate that spatio-temporal-spectral entrainment patterns allow groove ratings to be predicted with good accuracy, where ITPC provided more information than SBC.

## 4 Discussion

### 4.1 Summary

Previous research showed that the brain entrains to rhythmic acoustic stimuli such as music, but is was unclear whether and how neural entrainment to pop songs relates to feelings of their groove. Thus, the present study investigated how entrainment of neural oscillations in the delta-theta and beta bands to popular songs contributes to groove perception and enables prediction of this subjective psychological state. Our study combined and replicated stimuli and analyses procedures from Janata et al. [3] and Cameron et al. [38] to test if neural entrainment, as measured by EEG, can predict groove in popular music. Participants rated groove for 9 popular songs and 12 simple clapping rhythms while we collected EEG data. Univariate analyses revealed that music and clapping stimuli entrained the phase of neural oscillations in the delta and theta frequency bands, as measured by ITPC and SBC in fronto-central clusters. In these clusters, both ITPC and SBC of the songs stimuli in theta band in these channels correlated weakly, but significantly with groove ratings. Multivariate decoding analyses showed that groove ratings could be predicted from complex and distributed spatio-temporal-spectral patterns with high accuracy (up to *r* = 0.8), where ITPC was most informative for prediction.

### 4.2 Univariate Neural Entrainment

Our initial finding of significant entrainment of neural delta-theta oscillations by music and clapping stimuli was in line with previous studies reporting that rhythmic stimuli such as speech [21, 22], clapping rhythms [38], and simple music [23, 24, 25] entrain neural oscillations in these frequency bands in fronto-central channels. Notably, the profile of ITPC showed two peaks: The peak in the low delta band (around 1.3 Hz) presumably arose from entrainment to the beat of the stimuli [23, 7]. The second peak in the theta band (around 6 Hz) was even more pronounced and may be related to predictive processing [52], active auditory segmentation of the stimuli [53], or related to their melodic spectral complexity [25]. Analyses of entrainment over the range of channels and frequencies showed that ITPC measured even stronger and more consistent entrainment than SBC to the songs as well as clapping stimuli. Our univariate correlation analyses showed that both ITPC and SBC in fronto-central channels were weakly, but consistently and significantly correlated with groove ratings for song stimuli: Stronger entrainment was associated with higher groove ratings. We did not find significant neural entrainment or correlations with groove in the beta band, in contrast to previous studies that reported beta coherence as evidence of coupling between the auditory and motor sensory networks [32, 33]. Yet, the spatial specificity of the fronto-central cluster suggest that sensorimotor coupling in these regions may be critically involved in groove as previously assumed [27, 23]. Because our participants were instructed not to move along with the music in order to reduce EEG artifacts, the lack of evidence for beta synchronization may be the result of an inhibitory mechanism for motor activation (e.g., one that is guided by prefrontal control regions).

Our finding that theta entrainment predicts groove complements a recent MEG study which found that groove ratings of simple melodies can be decoded from oscillatory power in the delta band (at 2 Hz) from dorsal auditory pathways extending into motor regions [7]. However, control analyses of our data did not show that power and groove ratings were correlated for songs and clapping stimuli (see supplementary material Fig. 1). Importantly, univariate analyses did not find a significant correlation of ITPC and groove for our MIDI clapping stimuli, in contrast to the previous study reporting such a finding for physically performed clapping [38]. This null result likely arose from the restricted range of groove reported by the participants (see Fig. 2): Because participants consistently reported intermediate groove levels for all 12 clapping rhythms, these small variations could not be explained by ITPC or SBC. This result emphasizes that it is important to use stimuli that elicit a wide range of groove ratings when analyzing such brain-behavior relationships.

An important observation was that both measures of neural entrainment produced similar results, particularly with regard to their correlation with groove ratings. This is not self-evident, as even though ITPC and SBC both measure the synchronization of neural phase to stimulus dynamics, they capture different temporal aspects of entrainment that can diverge: ITPC measures the synchronization of the neural phase across stimulus replications at each time point of a stimulus presentation, while SBC measures moment-to-moment synchronization between the stimulus envelope and neural phase. Thus, ITPC emphasizes the consistency of neural phase across stimulus replications while neglecting the ongoing phase-relation to the phase dynamics of the stimulus. For example, ITPC could be high if the brain entrains to a stimulus with phase slips or resets compared to the stimulus, as long as the slips and resets are consistent across stimulus presentations. By contrast, SBC tracks the ongoing alignment of stimulus and neural phase while ignoring whether this alignment is consistent over multiple stimulus presentations. For example, SBC could be high if the brain entrains to a stimulus phase with a consistent phase relation for that stimulus, but this phase relation may differ for different stimulus presentations. In other words, the phase relationship could be positive in one trial and negative in another, resulting in a high SBC in both trials but a significant decrease in ITPC across trials.

Thus, our univariate analyses suggest that subjective groove is linked to consistent entrainment of a fronto-central mechanism to the temporal dynamics of the ongoing song, as well as a consistent entrainment over multiple replications of the song. This finding is in line with the predictive coding account of groove: The account assumes that entrainment in the theta band indexes a predictive process [52] which is linked to groove perception [29, 27]. If the brain predicts the upcoming rhythmical structure well despite some irregularity, the prediction mechanism leads to a consistent phase relation between song and neural phase, leading to strong neural entrainment and groove. Thus, different songs can elicit stronger or weaker groove ratings if predictions are consistently accurate in high-groove songs or more often fail in low-groove songs. Prediction errors, which may be captured by transient phase slips in neural data, reduce groove: Since the correlations between theta entrainment and groove were computed at the trial level, variations in entrainment were associated with variations in groove, even across different presentations of the same song. Therefore, moment-to-moment variations in prediction accuracy, as measured by theta entrainment, explain a significant proportion of the variability in subjective groove perception. Future research could examine music predictability [54, 55] and investigate an inverse U-shaped relationship between predictability and phase entrainment, akin to the syncopation-groove link [7].

### 4.3 Multivariate Decoding and Modeling

Our multivariate decoding approach used parametric and machine learning approaches like XGBoost and Random Forest to show that more complex spatio-temporal-spectral feature patterns of ITPC and SBC allow to predict trialwise groove ratings from neural entrainment up to correlations of 0.85. Still, although we found high within-participant rating consistency of individual songs and clapping stimuli, the decoding performance was much greater for the per-trial (individual) predictions than for the per-stimulus (averaged) predictions. This may be due to the participants slightly inconsistent rating behavior, which may arise from variations of neural entrainment between trials (see above), or the higher number of data points available for training and validating the models. Consistent with the univariate analyses, the decoding approach found much higher prediction accuracy for the songs as compared to clapping stimuli. As discussed above, the most likely reason is that clapping stimuli only led to a restricted range of groove ratings, thus limiting potential co-variation with neural entrainment. While univariate analyses revealed strongest correlations of SBC and groove, the multivariate decoding approach found highest prediction accuracy from distributed entrainment patterns measured with ITPC. Patterns of ITPC were much more informative than SBC because nearly as high correlations were achieved with ITPC patterns alone compared to the combination of SBC and ITPC. This apparent discrepancy of the decoding to the univariate approach could arise due to three reasons: First, the decoding approach exploited widely distributed information of entrainment patterns spanning temporal, spatial, and frequency dimensions that by far exceeded our narrower focus in univariate analyses. Inspection of the most predictive features revealed that at least for songs the most informative features came from fronto-central channels in the delta-theta band (Tab. 1 in the supplementary material), showing some consistency of the univariate and multivariate decoding approaches. However, the individual predictive power of even the most informative features was rather low (*r ≈* 0.3), suggesting that the information to predict groove was widely distributed and could not be attributed to a single source alone. This is in line with recent studies suggesting that neural activity in the delta-, theta- and beta-bands of distributed sensory-motor networks is involved in the perception of music and groove [23, 32, 33, 7, 34]. Second, the univariate analysis correlated neural entrainment with groove at an individual participant level. By contrast, the decoding approach involved training models using the entire group dataset, with the aim of creating a model that could be generalized and used to predict groove across all individuals. The decoding approach implicitly identified predictive patterns across all individuals (i.e., fixed-effects approach), while the univariate analyses tested the consistency of correlations under the assumption of variability between participants (i.e., randomeffects approach). Thus, the greater statistical power of the decoding approach [56] may have contributed to higher correlations. Third, it is important to keep the differential goals of the two approaches in mind: The univariate approach aimed at identifying and understanding neural mechanisms of groove, thus focusing on prespecified frequencies and channels. By contrast, the decoding approach aimed at improving prediction accuracy of groove from any complex neural entrainment patterns, at potential cost of explainability and transparency. While the univariate approach may help to refine neural models of groove [27, 29, 4], the decoding models can estimate groove objectively and accurately in a short-window, post-hoc manner (i.e. shortly after repeated listening to the stimuli). The decoding models can thus be applied for example to quantify how musical properties affect groove perception and overall music quality [4, 1, 43]. The high prediction accuracy of trial-wise entrainment data could be even leveraged to measure subjective groove perception online while listening to music. For example, closedloop technologies could be conceived that adapt music to the listener’s ongoing groove perception.

### 4.4 Limitations

Critically, several limitations of our study must be acknowledged: First, the neural source of the observed entrainment-groove correlation is still unclear. As we only used a rather low number of EEG channels, we did not attempt source localization. To possibly capture the neural origins of groove-related entrainment, future studies should investigate sensorimotor coupling to music stimuli in auditory and motor regions in source space, for which MEG provides better spatial resolution than EEG [7]. Second, differences in oscillatory power between songs and motor artifacts from head motion could have driven the correlation between entrainment and groove ratings. Regarding power confounds, we explicitly focused on phase-based entrainment measures which are in principle independent of power, but can be practically confounded with power if the stimuli elicited differential power [57]. However, control analyses did not show that power and groove ratings were correlated for songs and clapping stimuli (see supplementary material Fig. 1), which makes a power confound unlikely. To avoid head motion confounds, we explicitly instructed participants not to move their heads and to maintain central fixation on the screen. Yet, minor movements may still have happened unnoticed. However, we consider these confounds unlikely because typical motor responses to music synchronize to the beat or its sub-harmonics [58, 59]. Thus, head motion confounds should appear at frequencies at or below 2 Hz, which is a considerably lower frequency than our main findings around 6 Hz. Additionally, prohibiting movement may have forced participants to suppress a motor component which may be critical for sensorimotor coupling as a neural basis of groove, and may explain why we did not observe neural entrainment and correlation with groove in the beta band. Third, a relatively large proportion of trials were affected by artifacts (e.g., from eye-motion related artifacts, electrode drifts or jumps) due to their long duration. Because we wanted to keep the full dataset and assumed that these artifacts were not correlated to stimulus type or groove, we included all trials in our analyses. Therefore, noisy EEG data may have rendered the measurement of ITPC and SBC, as well as the correlation between neural entrainment and groove ratings, more unreliable. Fourth, we used clapping stimuli corresponding to the ones called *mechanical* in [38]. The correlation between our and the ones presented by Cameron et al. was low and negative (−0.20). While the difference in overall mean ratings across studies is small, relatively wide CIs in our data and likely similar CIs in their data (due to fewer ratings) make this relationship difficult to interpret and suggest it may primarily reflect statistical uncertainty. Additionally, approximately half of the participants reported struggling to provide meaningful ratings for these clapping stimuli. However, because we decoded ratings at the per-trial or per-stimulus level rather than based on mean ratings, we chose to retain the clapping stimuli. This may partially account for the comparatively low explanatory power observed for the models for the clapping datasets.

### 4.5 Future Work

In the future, we may need to focus on improved generalization by including a broader range of stimuli (genres, cultures, etc.) as well as a more diverse participant sample (country of origin, age, etc.). Based on this, cross-dataset validation might be possible and is likely needed. One can also examine additional behavioral markers of groove by not limiting participants’ movements. Using explainable AI and improved EEG systems with more channels, neurophysiological modeling can be further refined to improve understanding. Given that groove is assumed to have evolved primarily in group contexts, incorporating multi-listener and other social settings may also modulate groove ratings. Eventually, improving the practicality of the measurement can also be done by focusing on the most important EEG channels and therefore reducing the weight on a participant’s head and also the time in the experiments. Future studies may test which rhythmical features of the songs contribute to entrainment and groove perception [8, 15, 60], for example by systematically manipulating features such as syncopation. Furthermore, the causal role of delta-theta entrainment in groove sensations could be investigated by disturbing entrainment to music with additional external rhythmic auditory entrainment [61, 62], or with neurostimulation such as transcranial alternating current stimulation [63, 64]. Finally, decoding approaches like the ones presented here may be the basis of adaptive music applications or tools that adjust in response to a listener’s brain state, which can be of research, practical, and hedonic value.

### 4.6 Conclusion

Our study showed that both music and clapping stimuli consistently entrained neural oscillations in the delta-theta bands as measured by ITPC and SBC and that groove ratings can be modeled using such features. While univariate correlations revealed only weak but significant correlations between theta entrainment and groove ratings in fronto-central clusters for songs, multivariate decoding showed that spatio-temporal-spectral EEG patterns predicted groove significantly more accurately. Patterns of inter-trial phase coherence were found to be more informative than patterns of stimulus-brain coherence. Overall, our results support the notion that groove is rooted in the brain’s ability to entrain to ongoing dynamic temporal structures of music in the theta band.

## Author Contributions

T.R., D.K., and A.R. planned the experiment, D.K. performed the experiment, T.R. ran the EEG data processing, T.R. performed the univariate analysis, D.K. performed the multivariate analysis, T.R. and D.K. wrote the manuscript, and T.R., D.K., and A.R. edited the manuscript.

## Conflicts of Interest

The authors declare that there is no conflict of interest regarding the publication of this article.

## Acknowledgment

We thank Rakesh Rao Ramachandra Rao for his valuable feedback, careful review, and insightful questions that helped improve the quality and clarity of this work. We thank all volunteers for taking part in the tests and also our student assistants for their work in supervising such sessions. This work is part of the Interconnected Lab for MEdia Technology Analytics (ILMETA) project (number 438822823), which is funded by Deutsche Forschungsgemeinschaft (DFG).

## Footnotes

1 www.gtec.at/product/g-nautilus-research

## Notes

### Competing Interest Statement

The authors have declared no competing interest.

### Summary of Updates

Small edit to correct a statement in the Inter-Trial Phase Coherence results.

