## Supplemental Material for "Neural Entrainment in the Theta Band Predicts Groove Perception in Popular Music"

November 10, 2025

The supplementary material to the paper titled "Neural Entrainment in the Theta Band Predicts Groove Perception in Popular Music" is provided subsequently.

### 1 Decoding: Best Performing Univariate Correlations

The Tabs. 1 and 2 show the highest-performing features and coefficients for the songs and clapping dataset respectively.

### 2 Power

We performed control analyses that did not show that power and groove ratings were correlated for songs and clapping stimuli. See Fig. 1 for more details.

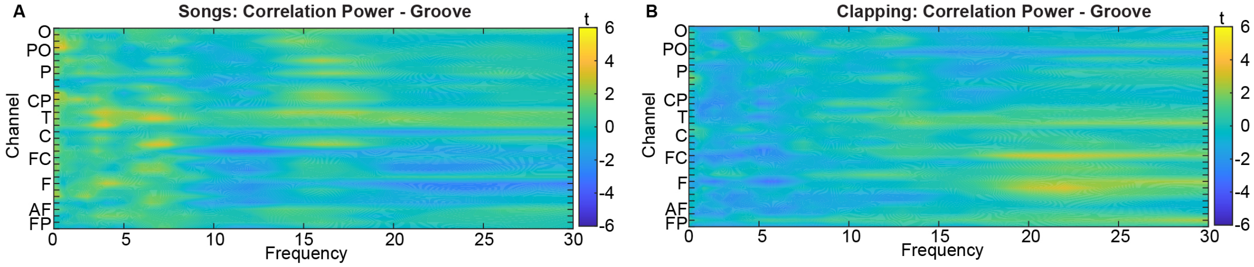


Figure 1: t-value maps of individual correlations between power and groove ratings be- tween 1-30 Hz in all 32 channels, for songs (A) and clapping (B). Electrode channels are ordered according to their posterior-anterior proximity (O = occipital; PO = parietal-occipital; P = parietal; CP = central-parietal; T = temporal; C = central; FC = frontal central; F = frontal; AF = anterior-frontal; FP = fronto-polar). Two-sided cluster-based corrected randomization t29 test did not identify significant clusters ($p > 0.05$)

| Data | Song Feature | n | PCC $r$ | SCC $\rho$ |
|---|---|---|---|---|
| aver. | SBC_freq_1.5_acroElec_sd | 270 | -0.330 | -0.313 |
| aver. | ITPC_FC1_freq_6.5_sd | 270 | 0.316 | 0.314 |
| ind. | ITPC_FC1_freq_6.5_sd | 810 | 0.299 | 0.295 |
| aver. | ITPC_FC1_freq_6.5_perc_90 | 270 | 0.292 | 0.278 |
| aver. | SBC_freq_1.5_acroElec_max | 270 | -0.292 | -0.293 |
| aver. | SBC_freq_7.0_acroElec_mean | 270 | 0.285 | 0.276 |
| aver. | SBC_freq_7.0_acroElec_max | 270 | 0.285 | 0.276 |
| aver. | ITPC_FC1_freq_6.0_perc_90 | 270 | 0.284 | 0.295 |
| aver. | ITPC_FP1_time_47_perc_75 | 270 | 0.283 | 0.256 |
| aver. | SBC_freq_7.0_acroElec_perc_90 | 270 | 0.280 | 0.281 |
| aver. | ITPC_FP1_freq_5.5_mean | 270 | 0.280 | 0.278 |
| ind. | ITPC_FC1_freq_6.5_perc_90 | 810 | 0.277 | 0.261 |
| aver. | ITPC_Pz_time_47_perc_75 | 270 | 0.276 | 0.284 |
| aver. | ITPC_FC1_freq_5.5_perc_75 | 270 | 0.276 | 0.287 |
| aver. | ITPC_FP1_freq_5.5_perc_75 | 270 | 0.274 | 0.278 |
| aver. | ITPC_FP1_freq_5.5_median | 270 | 0.274 | 0.270 |
| aver. | SBC_freq_7.0_acroElec_perc_75 | 270 | 0.273 | 0.275 |
| aver. | SBC_freq_7.0_acroElec_median | 270 | 0.272 | 0.250 |
| ind. | ITPC_FC1_freq_6.0_perc_90 | 810 | 0.269 | 0.278 |
| aver. | SBC_freq_6.5_acroElec_perc_90 | 270 | 0.268 | 0.255 |
| ind. | ITPC_FP1_time_47_perc_75 | 810 | 0.268 | 0.238 |
| ind. | ITPC_FP1_freq_5.5_mean | 810 | 0.265 | 0.261 |
| aver. | ITPC_FP1_freq_6.0_mean | 270 | 0.264 | 0.262 |
| aver. | SBC_FC2_freq_7.0 | 270 | 0.264 | 0.259 |
| aver. | ITPC_FC2_time_47_median | 270 | 0.262 | 0.240 |
| aver. | ITPC_FC1_freq_6.0_perc_75 | 270 | 0.262 | 0.254 |
| ind. | ITPC_Pz_time_47_perc_75 | 810 | 0.262 | 0.268 |
| ind. | ITPC_FC1_freq_5.5_perc_75 | 810 | 0.262 | 0.271 |
| aver. | SBC_freq_3.0_acroElec_sd | 270 | -0.262 | -0.257 |
| aver. | ITPC_FC1_freq_5.5_median | 270 | 0.261 | 0.263 |

Table 1: Best Pearson (PCC) and Spearman (SCC) correlation results of the *songs* dataset (top 30 features by absolute PCC). Features correlated with personal average groove rating per stimulus (aver.) or individual groove rating per trial (ind.).

| Data | Clapping Feature | n | PCC $r$ | SCC $\rho$ |
|---|---|---|---|---|
| aver. | SBC_freq_18.0_acroElec_sd | 360 | -0.249 | -0.188 |
| aver. | SBC_F3_all_perc_90 | 360 | -0.246 | -0.232 |
| aver. | SBC_freq_12.0_acroElec_sd | 360 | -0.244 | -0.151 |
| aver. | SBC_freq_7.0_acroElec_max | 360 | -0.243 | -0.139 |
| aver. | SBC_freq_18.0_acroElec_max | 360 | -0.242 | -0.176 |
| aver. | SBC_freq_12.0_acroElec_perc_10 | 360 | 0.241 | 0.146 |
| aver. | SBC_freq_20.0_acroElec_sd | 360 | -0.240 | -0.172 |
| aver. | SBC_freq_5.5_acroElec_sd | 360 | -0.238 | -0.133 |
| aver. | SBC_freq_13.0_acroElec_sd | 360 | -0.237 | -0.160 |
| aver. | SBC_freq_13.0_acroElec_max | 360 | -0.237 | -0.131 |
| aver. | SBC_freq_12.0_acroElec_max | 360 | -0.235 | -0.102 |
| aver. | SBC_freq_12.0_acroElec_min | 360 | 0.232 | 0.130 |
| aver. | SBC_freq_6.0_acroElec_min | 360 | 0.232 | 0.142 |
| aver. | SBC_freq_18.0_acroElec_perc_90 | 360 | -0.229 | -0.127 |
| aver. | SBC_freq_20.0_acroElec_min | 360 | 0.229 | 0.171 |
| aver. | SBC_freq_13.0_acroElec_perc_90 | 360 | -0.229 | -0.113 |
| aver. | SBC_freq_20.0_acroElec_max | 360 | -0.228 | -0.148 |
| aver. | SBC_freq_11.0_acroElec_sd | 360 | -0.228 | -0.178 |
| aver. | SBC_all_elec_all_freqs_perc_90 | 360 | -0.228 | -0.203 |
| aver. | SBC_freq_18.0_acroElec_min | 360 | 0.227 | 0.120 |
| aver. | SBC_freq_7.0_acroElec_sd | 360 | -0.226 | -0.132 |
| aver. | SBC_freq_12.0_acroElec_perc_90 | 360 | -0.226 | -0.099 |
| aver. | SBC_freq_13.0_acroElec_min | 360 | 0.224 | 0.144 |
| aver. | SBC_FC2_all_perc_90 | 360 | -0.223 | -0.218 |
| aver. | SBC_C3_all_perc_90 | 360 | -0.223 | -0.196 |
| aver. | SBC_freq_5.5_acroElec_min | 360 | 0.222 | 0.116 |
| aver. | SBC_freq_6.0_acroElec_sd | 360 | -0.222 | -0.131 |
| aver. | SBC_freq_24.0_acroElec_max | 360 | -0.221 | -0.192 |
| aver. | SBC_freq_6.5_acroElec_max | 360 | -0.220 | -0.147 |
| aver. | SBC_freq_13.0_acroElec_perc_75 | 360 | -0.217 | -0.106 |

Table 2: Best Pearson (PCC) and Spearman (SCC) correlation results of the *clapping* dataset (top 30 features by absolute PCC), all significant at $p < 0.001$ (***). Features correlated with personal average groove rating per stimulus.
